# Potentially highly potent drugs for 2019-nCoV

**DOI:** 10.1101/2020.02.05.936013

**Authors:** Duc Duy Nguyen, Kaifu Gao, Jiahui Chen, Rui Wang, Guo-Wei Wei

**Author notes:** Address correspondences to Guo-Wei Wei.

## Abstract

The World Health Organization (WHO) has declared the 2019 novel coronavirus (2019-nCoV) infection outbreak a global health emergency. Currently, there is no effective anti-2019-nCoV medication. The sequence identity of the 3CL proteases of 2019-nCoV and SARS is 96%, which provides a sound foundation for structural-based drug repositioning (SBDR). Based on a SARS 3CL protease X-ray crystal structure, we construct a 3D homology structure of 2019-nCoV 3CL protease. Based on this structure and existing experimental datasets for SARS 3CL protease inhibitors, we develop an SBDR model based on machine learning and mathematics to screen 1465 drugs in the DrugBank that have been approved by the U.S. Food and Drug Administration (FDA). We found that many FDA approved drugs are potentially highly potent to 2019-nCoV.

## 1 Introduction

The 2019 novel coronavirus (2019-nCoV) caused the pneumonia outbreak in Wuhan, China, in late December 2019 and has rapidly spread around the world. By Feb 5, 2020, more than 24000 individuals were infected and more than 490 fatalities had been reported. The World Health Organization (WHO) has declared this novel coronavirus outbreak a global health emergency. Currently, there is no specific antiviral drug for this epidemic. Considering the severity of this widespread dissemination and health threats, panic patients misled by media flocked to the pharmacies for Chinese Medicine herbs which were reported to “inhibit” the 2019-nCoV, despite no clinical evidence supporting the claim. Many researchers are engaged in developing anti-2019-nCoV drug s [1, 2]. However, new drug discovery and development is a long, costly and rigorous scientific process. A more effective approach is to search for anti-2019-nCoV therapies from the existing FDA-approved drug database.

Drug repositioning (also known as drug repurposing), which concerns the investigation of existing drugs for new therapeutic target indications, has emerged as a successful strategy for drug discovery due to the reduced costs and expedited approval procedures [3–5]. Several successful examples unveil its great values in practice: Nelfinavir, initially developed to treat the human immunodeficiency virus (HIV), is now being used for cancer treatments. Amantadine was firstly designed to treat influenza caused by type A influenza viral infection and is being used for Parkinson’s disease later on [6]. In recent years, the rapid growth of drug-related datasets, as well as open data initiatives, has led to new developments for computational drug repositioning, particularly, structural-based drug repositioning (SBDR). Machine learning, network analysis, and text mining and semantic inference are three major computational approaches commonly applied in drug repositioning [7]. The rapid accumulation of genetic and structural databases [8], the development of low-dimensional mathematical representations of complex biomolecular structures [9, 10], and the availability of advanced deep learning algorithms have made machine learning-based drug reposition a promising approach [7]. Considering the urgent need for anti-2019-nCoV drugs, a computational drug repositioning is one of the most feasible strategies for discovering 2019-nCoV drugs.

In SBDR, one needs to select one or a few effective targets. Study shows that 2019-nCoV genome is very close to that of the severe acute respiratory syndrome (SARS)-CoV [11]. The sequence identities of 2019-nCoV 3CL protease, RNA polymerase, and the spike protein with corresponding SARS-CoV proteins are 96.08%, 96%, and 76%, respectively [12]. We, therefore, hypothesize that a potent SARS 3CL protease inhibitor is also a potent 2019-nCoV 3CL protease inhibitor. Unfortunately, there is no effective SARS therapy at present. Nevertheless, the X-ray crystal structure of SARS 3CL protease has been reported [13] and the binding affinities of 115 potential SARS 3CL protease inhibitors are available in ChEMBL database [14]. Additionally, there are 15,843 protein-ligand complexes in PDBbind 2018 general set with binding affinities and X-ray crystal structures [15]. Moreover, the DrugBank contains about 1600 drugs approved by the U.S. Food and Drug Administration (FDA) [16]. The aforementioned information provides a sound foundation to develop an SBDR machine learning model for 2019-nCoV 3CL protease inhibition.

Recently, we have developed low-dimensional mathematical representations [9, 10] to reduce the structural complexity of macromolecules based on abstract mathematics, such as algebraic topology [17–20], differential geometry, and spectral graph theory [10, 21]. We exploit these representations to extract critical chemical and biological information for protein-ligand pose selection, binding affinity ranking, prediction, ranking, scoring, and screening [9, 10]. Paired with various machine learning, including deep algorithms, these approaches are the top competitor for D3R Grand Challenges, a worldwide competition series in computer-aided drug design in the past few years [22, 23].

In responding to the pressing need for anti-2019-nCoV medications, we develop mathematics-based deep learning models to systematically eventuate FDA approved drugs in the DrugBank for 2019-nCoV 3CL protease inhibition. With the consensus of two deep learning models based on convolutional neural networks and multitask deep learning, we report the top 15 potentially highly potent anti-2019-nCoV 3CL inhibitors, which provide timely guidance for the further development of anti-2019nCoV drugs.

## 2 Results

### 2.1 Sequence identity analysis

The sequence identity is defined as the percentage of characters that match exactly between two different sequences. The sequence identities between 2019-nCoV protease and the protease of SARS-CoV, MERS-CoV, HKU-1, OC43, HCoVNL63, 229E, and HIV are 96.1%, 52.0%, 49.0%, 48.4%, 45.2%, 41.9%, and 23.7%, respectively. It is seen that 2019-nCoV protease is very close to SARS-CoV protease, but is distinguished from other proteases. Clearly, 2019-nCoV has a strong genetic relationship with SARS-CoV, the sequence alignment in Figure 1 further confirms their relationship. Additionally, the available experimental data of SARS-CoV protease inhibitors can be used as the training set to generate new inhibitors of 2019-nCoV protease.

**Figure 1:**
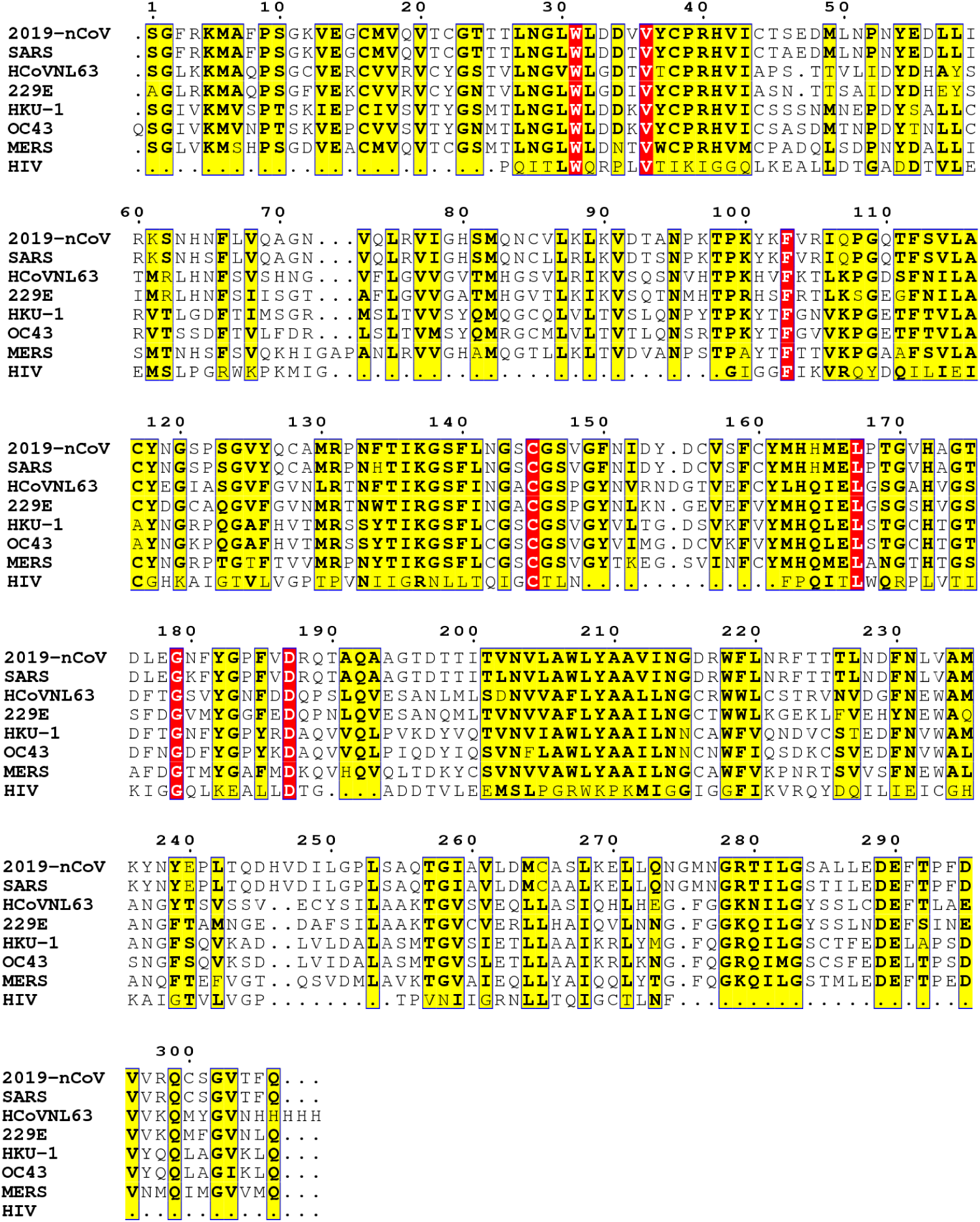
The protease sequence alignment between 2019-nCoV, SARS, MERS, OC43, HCoVNL63, HKU-1, 229E, and HIV.

### 2.2 Structure similarity analysis

Since the sequences are highly identical, the 2019-nCoV protease structure can be built by homology modeling with the SARS-CoV 3CL protease (PDB ID: 2A5I) [13] as a template. It turns out, as shown in Fig. 2, the homology structure of the 2019-nCoV protease is essentially identical to the X-ray structure of SARS-CoV 3CL protease. Particularly, the RMSD of two structures at the binding site is 0.21 Å. The high structural similarity between the two proteases suggests that anti-SARS-CoV chemicals can be equally effective for the treatment of 2019-nCoV.

**Figure 2:**
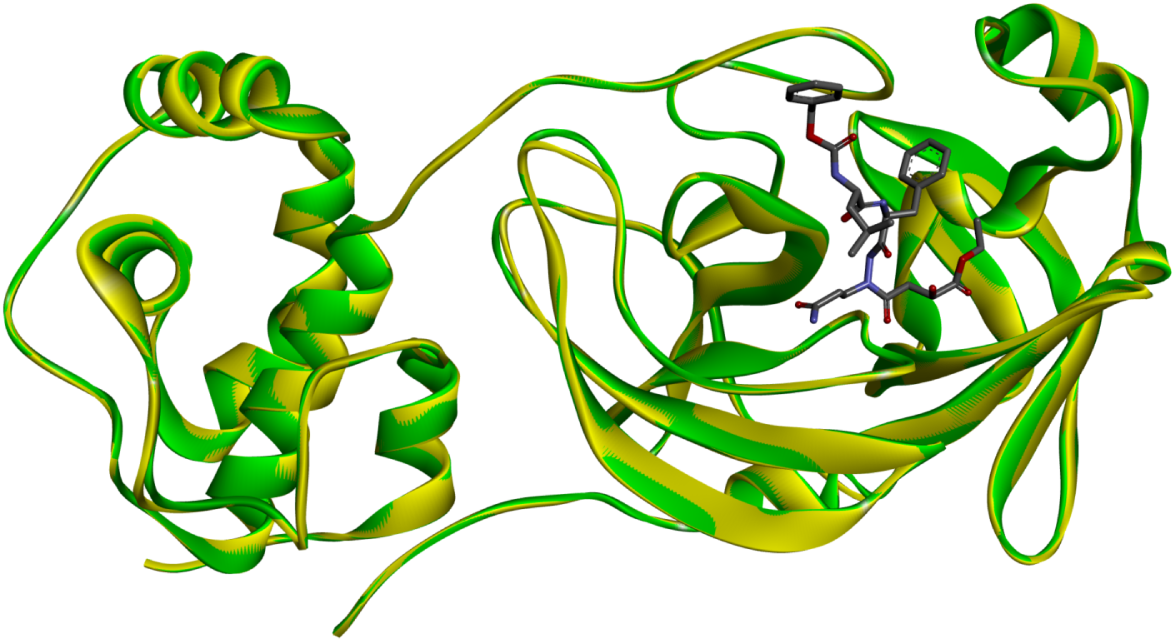
Illustration of the similarity and difference between protease structures of 2019-nCoV 3CL protease (in gold) and SARS-CoV 3CL protease (PDB ID: 2A5I, in green). The anti-SARS inhibitor in dark color indicates the binding site.

### 2.3 Binding analysis

We predict the binding affinities of 1465 3D FDA-approved drugs and 2019-nCoV protease complexes using two models, 3DALL and 3DMT. 3DALL is built with deep convolutional neural networks (CNNs) using the algebraic topology-based representation of protein-ligand complexes, with 84 SARS-CoV protease inhibitors and 15843 complexes from the PDBbind 2018 general set as the training set. 3DMT, a deep multitask CNN model based on the algebraic topology representation of protein-ligand complexes is the second model. In the current work, two tasks were developed in the 3DMT. The first task involves 1465 2019-nCoV protease complexes as the test set and 84 SARS-CoV protease inhibitors as the training set. The second task is trained with the PDBbind 2018 general set of 15843 protein-ligand complexes. Our top 15 potential 2019-nCoV inhibitors based on consensus binding affinities of the aforementioned two models are listed in Table 1. A complete list of predicted binding affinities of 1465 FDA-approved drugs is given in the Supplementary Material.

**Table 1:** A summary of potentially highly potent anti2019-nCoV drugs with predicted binding free energies (unit: kcal/mol) and corresponding trade names.

| DrugID | Type | Trade Name | Predicted Binding Energy |
| --- | --- | --- | --- |
| DB00188 | Bortezomib | Velcade, Chemobort, Bortecad | -12.15 |
| DB00690 | Flurazepam | Dalmane, Dalmadorm, Fluzepam | -10.38 |
| DB08901 | Ponatinib | Iclusig | -10.25 |
| DB00398 | Sorafenib | Nexavar | -10.01 |
| DB01254 | Dasatinib | Sprycel, Dasanix | -9.87 |
| DB01384 | Paramethasone | Cortidene Depot, Dilar, Dilarmine | -9.71 |
| DB00838 | Clocortolone | Victrelis | -9.58 |
| DB00301 | Flucloxacillin | Flora, Flox, Floxapen | -9.57 |
| DB06144 | Sertindole | Serdolect and Serlect | -9.54 |
| DB04920 | Clevidipine | Cleviprex | -9.52 |
| DB00673 | Aprepitant | Emend | -9.49 |
| DB01076 | Atorvastatin | Lipitor, Sortis | -9.49 |
| DB01594 | Cinolazepam | Gerodorm | -9.47 |
| DB00845 | Clofazimine | Lamprene | -9.43 |
| DB06717 | Fosaprepitant | Emend, Ivemend | -9.39 |

We briefly describe the predicted potentially highly potent anti-2019-nCoV drugs. The most potent one is Bortezomib, an anti-cancer medication, which is known as proteasome inhibitor and can be used to treat multiple myeloma and mantle cell lymphoma. The second drug is Flurazepam, which is a benzodiazepine derivative that possesses anxiolytic, anticonvulsant, hypnotic, sedative, and skeletal muscle relaxant properties. The third one, Ponatinib, an oral drug for the treatment of chronic myeloid leukemia and Philadelphia chromosome-positive acute lymphoblastic leukemia, which is a multi-targeted tyrosine kinase inhibitor. It is important to notice that this drug has the risk of life-threatening blood clots and severe narrowing of blood vessels. The next one is Sorafenib, a kinase inhibitor for the treatment of primary kidney cancer and liver cancer. The fifth drug, Dasatinib, is a therapy for treating certain cases of chronic myelogenous leukemia and acute lymphoblastic leukemia. The next one, Paramethasone, is a glucocorticoid with the general properties of corticosteroids. The seventh drug is Clocortolone, a topical steroid that is used in the form of an ester, clocortolone pivalate. It is interesting to note that this drug is always applied as a cream for the treatment of dermatitis. It is considered a medium-strength corticosteroid. Therefore, this drug might be used to clean 2019-nCoV contaminated materials, offering an extra layer of protection. The number eight drug, Flucloxacillin, is a narrow-spectrum beta-lactam antibiotic of the penicillin class. It is used to treat infections caused by susceptible Gram-positive bacteria. The next one, Sertindole, is an antipsychotic medication. The number ten drug, Clevidipine, is a dihydropyridine calcium channel blocker that used for the reduction of blood pressure when oral therapy is not feasible or not desirable. The eleventh drug, Aprepitant, is used to prevent chemotherapy-induced nausea and vomiting, as well as postoperative nausea and vomiting. The number twelve, Atorvastatin, is a statin drug used to prevent cardiovascular disease in those at high risk and treat abnormal lipid levels. The next drug is Cinolazepam, a benzodiazepine derivative. It possesses anxiolytic, anticonvulsant, sedative, and skeletal muscle relaxant properties. The number fourteen drug, Clofazimine, is used together with rifampicin and dapsone to treat leprosy. The fifteenth number drug, Fosaprepitant, is an antiemetic medication used in the prevention of acute and delayed nausea and vomiting associated with chemotherapy treatment.

## 3 Discussion

### 3.1 The structural analysis of top 3 potent drug candidates

The top-ranking candidate of the existing drugs is Bortezomib (see Figure 3(b)). Its predicted binding affinity to the nCoV-2019 protease is -12.29 kcal/mol. The high binding affinity is due to the strong hydrogen bond network formed between the drug and the nCoV-2019 protease. For example, the strongest hydrogen bonds are formed by two O atoms in two hydroxyls on the head of Bortezomib and three different aminos in the main chains of residues Gly143, Ser144, and Cys145 of nCoV-2019 protease. Therefore, the head bonds tightly with the side chains of the aforementioned residues. The other two important hydrogen bonds are located at the tail of the drug molecule. The first one is between the O atom in the Hydroxyl on the tail and the two H atoms in the amino acid of the main chain of Glu166 and the methyl of the main chain of Met165. The second one is the H atom in the amino on the tail and the O atom in the side chain of Gln189. As a result, the head, body, and tail of Bortezomib interact firmly with the protease binding site.

**Figure 3:**
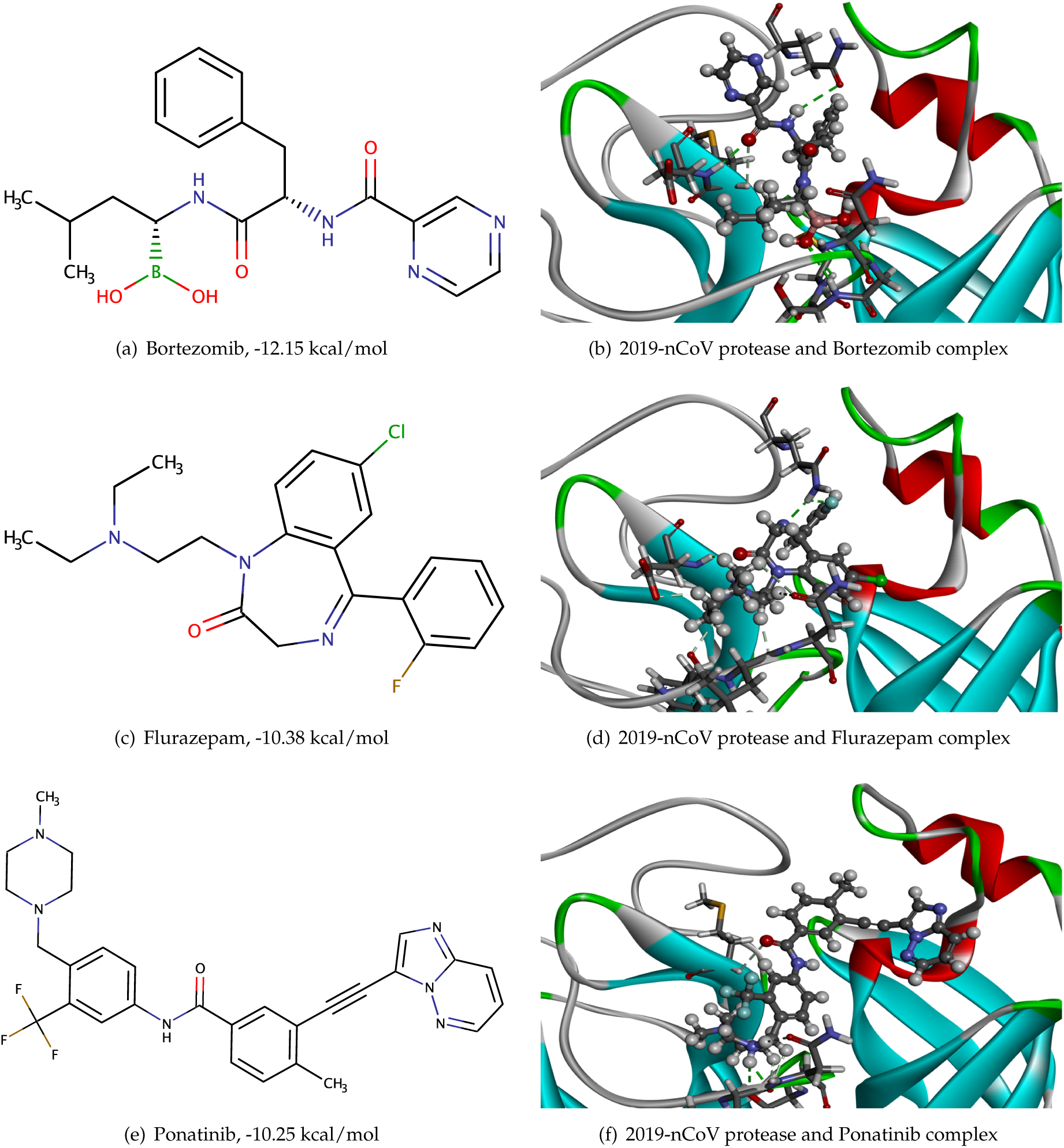
Bortezomib, Flurazepam, Ponatinib and their complexes with 2019-nCoV protease.

The second-best drug is Flurazepam (see Figure 3(d)) with a binding affinity of -10.37 kcal/mol. The strong hydrogen bonds between this molecule and the protease are formed by five different H atoms on the head of the drug with four different O atoms in the main chains of Phe140, Leu141, as well as the side chains of Asn142 and Glu166. Another important bond is formed by the H atom in the amino of the side chain of Gln189 with the F atom of the fluorobenzene and one N atom of the 1,4-diazepane in the drug. Additionally, the O atom in the drug adjacent to the 1,4-diazepane is bonded with the amino H atom of the side chain of Glu166. Therefore, the head, tail, and body of the molecule are firmly fixed to the binding site, which promises a strong binding to the 2019-nCoV protease.

The third one, Ponatinib (see Figure 3(f)), has a binding affinity -10.29 kcal/mol. The strong hydrogen bonds between this molecule and the protease are formed by two H atoms of the piperazine with the O atom in the side chain of Ser144 and the main chain of Leu141. Additionally, a bond exists between the O atom in the main chain of the drug and the H atom in the methyl of the main chain of Met165. These hydrogen bonds lead to a high binding affinity with 2019-nCoV protease.

The 3D complexes of 2019-nCoV protease and other 12 potential drugs are given in Supplementary Material.

### 3.2 Binding affinities of anti-virus protease drugs

It is interesting to analyze the predicted binding affinities of existing antiviral drugs developed as protease inhibitors. Their binding affinities are listed in Table 2. It is interesting to see that except for Boceprevir, which is a protease inhibitor used to treat hepatitis caused by the hepatitis C virus (HCV), the rest of protease inhibitors do not have a strong effect on 2019-nCoV. The predicted values by a recent study [24] are given in the parenthesis. It appears that these values are overestimated.

**Table 2:** A summary of predicted binding affinities (unit: kcal/mol) of antiviral protease inhibitors. Numbers in parenthesis are results from the literature [24].

| DrugID | Predicted Binding Energy | DrugID | Predicted Binding Energy |
| --- | --- | --- | --- |
| Boceprevir | -9.36 | Ritonavir | -7.19 (-8.47) |
| Tipranavir | -8.87 | Lisinopril | -7.17 |
| Dabigatran etexilate | -8.23 | Enalapril | -7.15 |
| Rivaroxaban | -7.88 | Vildagliptin | -7.15 |
| Fosamprenavir | -7.82 | Lopinavir | -7.12 |
| Argatroban | -7.81 | Apixaban | -7.09 |
| Sitagliptin | -7.79 | Perindopril | -7.06 |
| Saquinavir | -7.75 | Darunavir | -7.05 |
| Candoxatril | -7.62 | Ecabet | -6.86 |
| Simeprevir | -7.52 (-8.29) | Cilastatin | -6.86 |
| Telaprevir | -7.50 | Cilazapril | -6.85 |
| Saxagliptin | -7.49 | Quinapril | -6.80 |
| Indinavir | -7.46 | Nelfinavir | -6.74 |
| Linagliptin | -7.33 | Amprenavir | -6.73 |
| Atazanavir | -7.28 (-9.57) | Moexipril | -6.65 |
| Ramipril | -7.28 | Spirapril | -6.63 |
| Fosinopril | -7.28 | Trandolapril | -6.61 |
| Ximelagatran | -7.27 | Benazepril | -6.43 |
| Alogliptin | -7.26 | Captopril | -6.05 |
| Remikiren | -7.21 | Isoflurophate | -4.94 |

## 4 Material and methods

Our deep learning-based drug repositioning models employ mathematical pose (MathPose) and mathematical deep learning (MathDL) to predict 3D poses and protein-ligand binding affinities. The latter is used as a major criterion for searching anti-2019-nCoV therapies from the existing FDA-approved drugs. We first build a 3D 2019-nCoV 3CL protease structure by using homology modeling. A set of SARS-CoV protease inhibitors are docked to the 3D 2019-nCoV 3CL protease structure using our MathPose. The resulting complexes are used as a set of machine learning training. Additionally, a set of protein-ligand complexes from the PDBBind database is collected as another machine learning training set. Our training accuracy in terms of the Pearson correlation coefficient is higher than 0.99 in all deep learning models.

### 4.1 3D 2019-nCoV protease structure

Homology modeling, a procedure that constructs an atomic-resolution model of a protein from its amino acid sequence and experimental 3D structure of the related homologous protein, i.e., the “template,” is used to generate the 3D structure of 2019-nCoV 3CL protease. The SWISS model (https://swissmodel.expasy.org/) is employed with the protease structure of SARS-CoV (PDB ID: 2A5I [13]) as a template. The sequence identity between the 3CL proteases of SARS-CoV and 2019-nCoV is 96.08%.

### 4.2 SARS-CoV protease inhibitor dataset

ChEMBL [14], an open database that brings chemical, bioactivity, and genomic data together to translate genomic information into effective new drugs, is employed to construct our 2019-nCoV training set. Considering the high sequence identity between viral proteases of 2019-nCoV and SARS-CoV, we take the protease of SARS-CoV as the input target in ChEMBL and a total 115 ChEMBL IDs of the target can be found. The experimental Δ*G* values of 2019-nCoV 115 SARS-CoV protease inhibition compounds range from −10.0 kcal/mol to 7.5 kcal/mol. We exclude compounds with positive values, resulting in a total of 84 SARS-nCoV protease inhibition compounds for our machine learning training. A collection of these 84 compounds is given in the Supplementary Materials.

### 4.3 Binding affinity training set

The PDBbind database is a yearly updated collection of experimentally measured binding affinity data (Kd, Ki, and IC50) for the protein-ligand complexes deposited in the Protein Data Bank (PDB). The PDBbind general set, instead of the high-quality refined set, is chosen as our training set because of the FDA approved drugs involve a wide range of protein targets. In the current work, we use a set of 15,843 X-ray crystal structures of protein-ligand complexes and associated binding affinities from the PDBbind v2018 general set [15]. The information of these complexes is provided in the Supplementary Materials.

### 4.4 FDA approved drugs

DrugBank (www.drugbank.ca) is a richly annotated, freely accessible online database that integrates massive drug, drug target, drug action, and drug interaction information about FDA-approved drugs with the experimental drugs which are going through the FDA approval process [16]. Due to the high quality and sufficient information contained in, the DrugBank has become one of the most popular reference drug resources used all over the world. A total of 1553 FDA-approved drugs are contained in the DrugBank. However, in the present work, a number of FDA-approved drugs encountered difficulties in docking with the target molecule. Therefore, the MathPose successfully created 3D protein-ligand complex structures for 1465 FDA-approved drugs and 2019-nCoV protease.

### 4.5 MathDL

MathDL, designed for predicting various druggable properties of 3D molecules [23], is capable of efficiently and accurately encoding the high-dimensional biomolecular interactions into low-dimensional representations. Algebraic graph theory-based algorithms [25], differential geometry, and algebraic topology methods [23] are applied to generate three mathematical representations of data in MathDL. These data representations can be integrated with well-designed deep learning models, such as gradient-boosted trees (GBTs) and convolutional neural networks (CNNs), for pose ranking and binding affinity predictions. In D3R Grand Challenges (https://drugdesigndata.org/about/grand-challenge), a worldwide competition series in computer-aided drug design, MathDL had been proved as the top performer in free energy prediction and ranking [22, 23]. Figure 4 illustrates the framework of the MathDL model, which combined the aforementioned mathematical representations with the CNN architecture for druggable properties predictions. The PDBbind 2018 general set [15], along with the SARS 3CL protease related dataset is used in our training process. In this section, we briefly describe the algebraic topology representation used in the present work. Details can be found in the literature [23].

**Figure 4:**
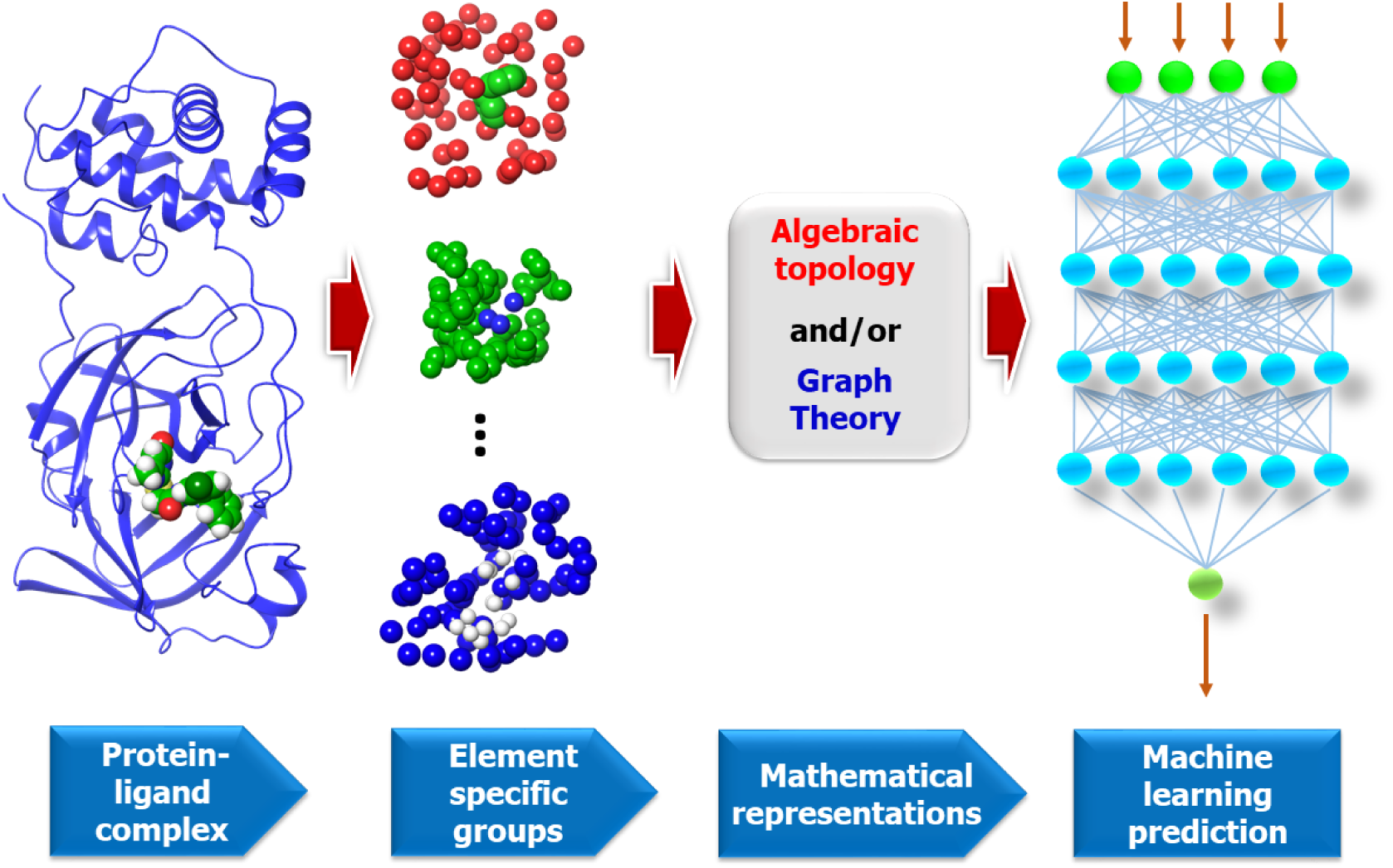
A framework of MathDL energy prediction model which integrates advanced mathematical representations with sophisticated CNN architectures

#### 4.5.1 Algebraic topology-based representation

Even with a glimpse of topology, one can realize it dramatically simplifies geometric complexity [9, 17– 20]. The study of topology reveals characterizes of different dimensions. As a type of algebraic topology, simplicial homology studies complexes on discrete datasets under various settings, such as the Vietoris-Rips (VR) complex, Čech complex or alpha complex, and identifies the topological invariants of a point-cloud dataset such as atomic coordinates in a protein [26]. Separated components, rings, and cavities can be classified for a given configuration and their numbers are referred to as Betti-0, Betti-1, and Betti-2, respectively. In this topological analysis process, the metrics or coordinates are fully abandoned. Instead, geometric and topological information is captured as data representation. Moreover, as a new development branch of algebraic topology, persistent homology which combines multiscale geometric information and topological invariants to achieve a geometry-enriched topological characteristic, e.g., barcodes. Therefore, the “birth” and “death” of separated components, circles, rings, voids or cavities can be indicated at all spatial scales by topological measurements. Key concepts are briefly shown as following.

In algebraic topology, simplices are the essential building blocks. Let *v*_0_, *v*_1_, *v*_2_, …, *v*_*k*_ be *k* +1 affinely independent points. A (geometric) *k*-simplex *σ*^*k*^ is the linear combinations of these points in ℝ^*n*^ (*n* ≥ *k*), whose coefficients are positive and satisfy that their summation equals to 1. For example, a 0, 1, 2, or 3-simplex is considered as a vertex, an edge, a triangle, or a tetrahedron, respectively. A simplicial complex *K* is a topological space composed of simplices which satisfies that every face of a simplex *σ*_*k*_ ∈ *K* is also in *K* and the non-empty intersection of any two simplices is a face for both. To identify the homology group, a *k*-chain [*σ*^*k*^] is a summation 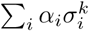 of *k*-simplices 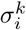, and the set of all *k*-chains of the simplicial complex *K* equipped with an algebraic field (typically, ℤ_2_) forms an abelian group *C*_*k*_(*K*, ℤ_2_). The homology defined on a series of abelian groups is used to analyze topological invariants which requires boundary operators to connect these chain spaces. The boundary operators *∂*_*k*_ : *C*_*k*_ → *C*_*k*−1_ for a *k*-simplex *σ*^*k*^ = {*v*_0_, *v*_1_, *v*_2_, …, *v*_*k*_} are homomorphisms defined as 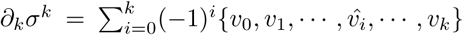, where 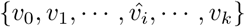 is a (*k* − 1)-simplex excluding *v*_*i*_ from the vertex set. Consequently, a important property of boundary operator, *∂*_*k*−1_*∂*_*k*_ = ∅, follows from that boundaries are boundaryless. The algebraic construction to connect a sequence of complexes by boundary maps is called a chain complex

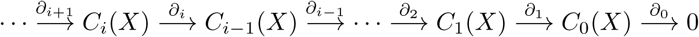

and the *k*th homology group is the quotient group defined by

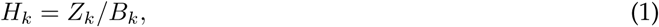

where the *k*-cycle group *Z*_*k*_ and the *k*-boundary group *B*_*k*_ are the subgroups of *C*_*k*_ defined as, *Z*_*k*_ = ker*∂*_*k*_ = {*c* ∈ *C*_*k*_ | *∂*_*k*_*c* = ∅}, *B*_*k*_ = im *∂*_*k*+1_ = {*∂*_*k*+1_*c* | *c* ∈ *C*_*k*+1_}. The aforementioned property implies *B*_*k*_ ⊆ *Z*_*k*_ ⊆ *C*_*k*_. The Betti numbers are defined by the ranks of *k*th homology group *H*_*k*_ which counts *k*-dimensional holes, especially, *β*_0_ = rank(*H*_0_) reflects the number of connected components, *β*_1_ = rank(*H*_1_) reflects the number of loops, and *β*_2_ = rank(*H*_2_) reveals the number of voids or cavities. Together, the set of Betti numbers {*β*_0_, *β*_1_, *β*_2_, …} indicates the intrinsic topological property of a system.

Persistent homology [18] is devised to track the multiscale topological information over different scales along a filtration. A filtration of a topology space *K* is a nested sequence of subspaces {*K*^*t*^}_*t*=0,…,*m*_ of *K* such that ∅ = *K*^0^ ⊆ *K*^1^ ⊆ *K*^2^ ⊆ … ⊆ *K*^*m*^ = *K*. Moreover, on this complex sequence, we obtain a sequence of chain complexes by homomorphisms: *C*_*_(*K*^0^) → *C*_*_(*K*^1^) → … → *C*_*_(*K*^*m*^) and a homology sequence: *H*_*_(*K*^0^) → *H*_*_(*K*^1^) → … → *H*_*_(*K*^*m*^), correspondingly. The *p*-persistent *k*th homology group of *K*^*t*^ is defined as

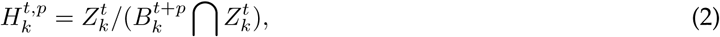

where 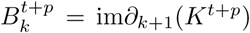. Intuitively, this homology group records the homology classes of *K*^*t*^ that are persistent at least until *K*^*t*+*p*^. Under the filtration process, the persistent homology barcodes can be generated. To make use of advanced deep learning algorithms, we vectorize persistent homology barcodes by dividing them into bins and calculating persistence, birth, and death incidents in each bin. Furthermore, the statistics of element-specific persistent homology barcodes are taken into consideration as well in fixed-length features.

### 4.6 MathPose

MathPose, a 3D pose predictor which converts SMILES strings into 3D poses with references of target molecules, was the top performer in D3R Grand Challenge 4 in predicting the poses of 24 beta-secretase 1 (BACE) binders [23]. For one SMILES string, around 1000 3D structures can be generated by a common docking software tool, i.e., GLIDE [27]. Moreover, a selected set of known complexes is re-docked by the three docking software packages mentioned above to generate at 100 decoy complexes per input ligand as a machine learning training set. The machine learning labels will be the calculated root mean squared deviations (RMSDs) between the decoy and native structures for this training set. Furthermore, MathDL models will be set up and applied to select the top-ranked pose for the given ligand. Additionally, the top poses will be fed into the MathDL for druggable proprieties evaluation.

## 5 Conclusion

The current pneumonia outbreak caused by a new coronavirus (CoV), called 2019-nCoV in China, has evolved into a global health emergency declared by the World Health Organization. Although there is no effective anti-viral medicine for the 2019-nCoV, the 3CL proteases of 2019-nCoV and SARS-CoV have a sequence identity of 96%, which provides a foundation for us to hypothesize that all potential anti-SARS-CoV chemotherapies are also effective anti-2019-CoV molecules. We build a three-dimensional (3D) 2019-nCoV 3CL protease structure using a SARS-CoV 3CL protease crystal structure as a template and collect a set of 84 SARS-CoV inhibition experimental data. The molecules of this set are docked to the 3D 2019-nCoV 3CL protease structure to form a machine learning training set. Additionally, the PDBbind 2018 general set of 15,843 protein-ligand complexes is also included as an additional machine learning training set. Using these training sets, we develop two deep learning models based on low-dimensional algebraic topology representations of macromolecular complexes. A total of 1465 FDA-approved drugs is evaluated by their binding affinities predicted by the consensus of two models built with 1) a combination of algebraic topology and deep convolutional neural networks (CNNs), and 2) a combination of algebraic topology and deep multitask CNNs. According to the predicted binding affinities, we recommend many FDA-approved drugs as potentially highly potent medications to 2019-nCoV, which serve as a crucial step for the development of anti-2019-nCoV drugs.

## Supplementary Materials

Supplementary Materials are available online for 3D structure information and affinities of SARS-CoV inhibitors, FDA-approved drugs, and PDBbind data set.

## Acknowledgments

This work was supported in part by NIH grant GM126189, NSF Grants DMS-1721024, DMS-1761320, and IIS1900473, Bristol-Myers Squibb, and Pfizer.

