## Supplementary Material for "Potentially highly potent drugs for 2019-nCoV"

### Supplementary Materials
### Potentially highly potent drugs for 2019-nCoV

Duc Duy Nguyen[1], Kaifu Gao[1], Jiahui Chen[1], Rui Wang[1], and Guo-Wei Wei[1,2,3] *
[1] Department of Mathematics, Michigan State University, MI 48824, USA
[2] Department of Biochemistry and Molecular Biology
Michigan State University, MI 48824, USA
[3] Department of Electrical and Computer Engineering
Michigan State University, MI 48824, USA

### S1 Supplementary Figures

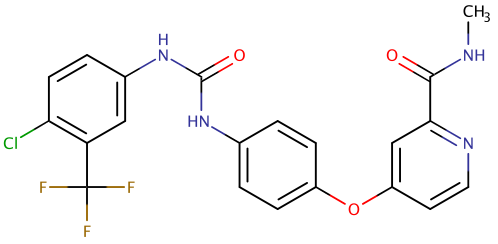

(a) Sorafenib, -10.01 kcal/mol

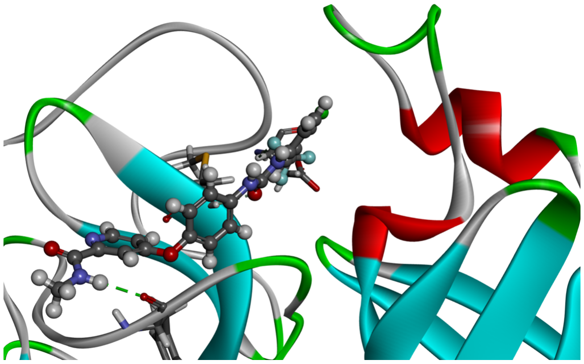

(b) 2019-nCoV protease and Sorafenib complex

Figure 1: Sorafenib and its complex with 2019-nCoV protease.

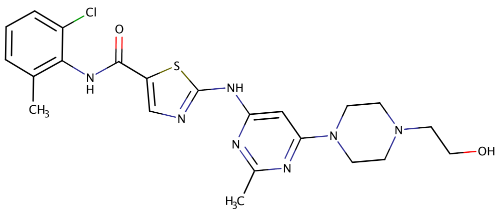

(a) Dasatinib, -9.87 kcal/mol

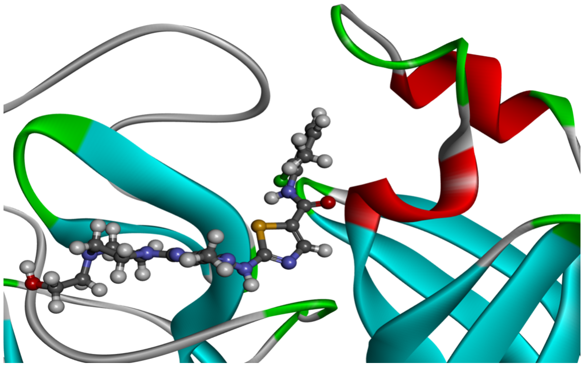

(b) 2019-nCoV protease and Dasatinib complex

Figure 2: Dasatinib and its complex with 2019-nCoV protease.

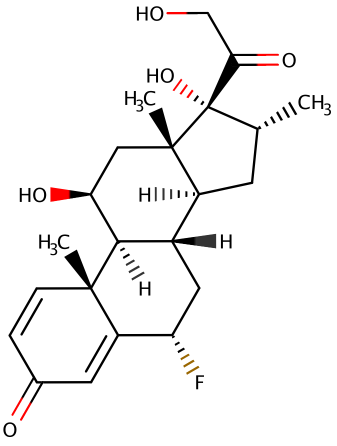

(a) Paramethasone, -9.71 kcal/mol

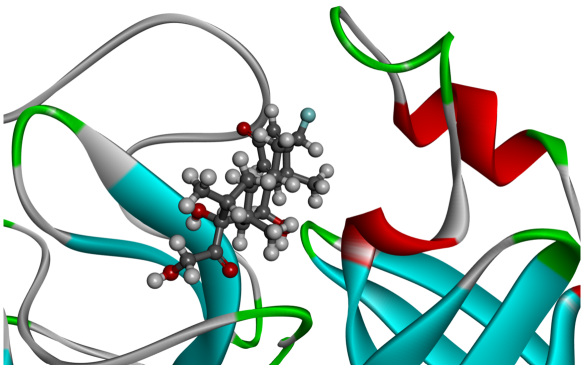

(b) 2019-nCoV protease and Paramethasone complex

Figure 3: Paramethasone and its complex with 2019-nCoV protease.

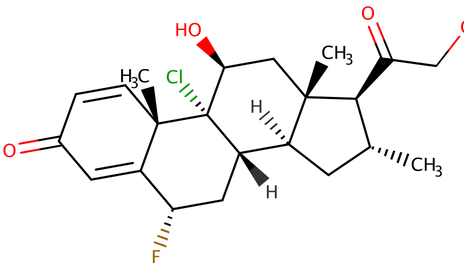

(a) Clocortolone, -9.58 kcal/mol

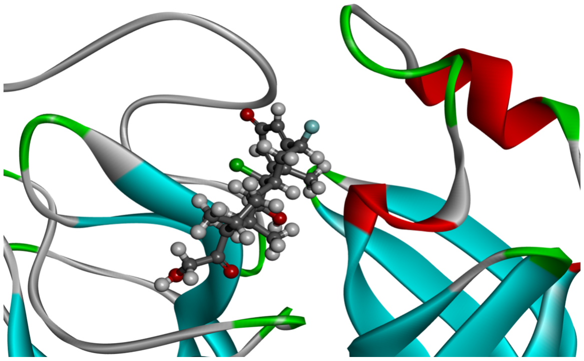

(b) 2019-nCoV protease and Clocortolone complex

Figure 4: Clocortolone and its complex with 2019-nCoV protease.

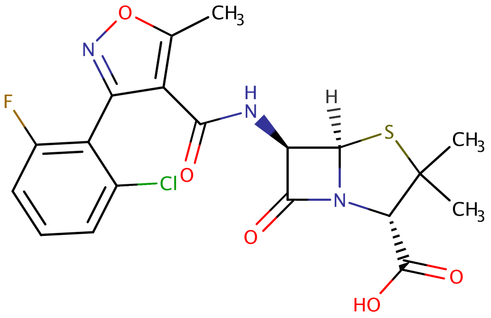

(a) Flucloxacillin, -9.57 kcal/mol

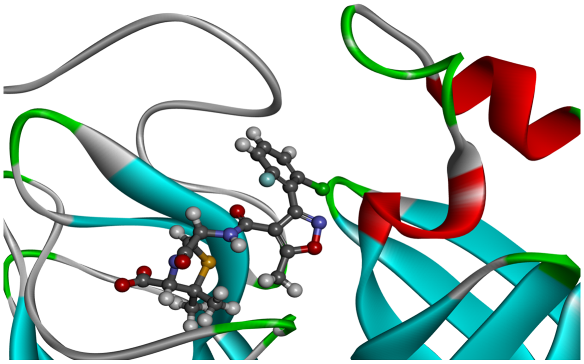

(b) 2019-nCoV protease and Flucloxacillin complex

Figure 5: Flucloxacillin and its complex with 2019-nCoV protease.

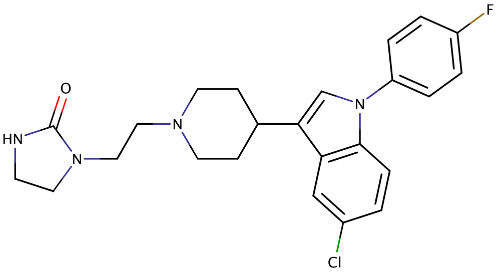

(a) Sertindole, -9.54 kcal/mol

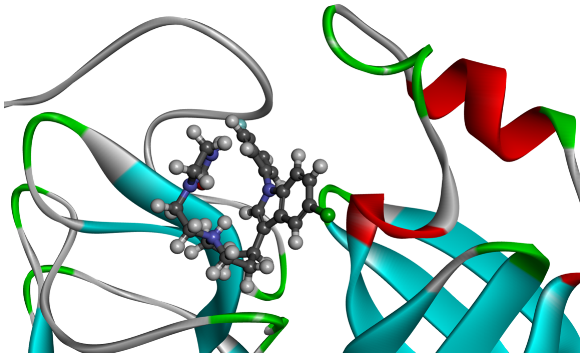

(b) 2019-nCoV protease and Sertindole complex

Figure 6: Sertindole and its complex with 2019-nCoV protease.

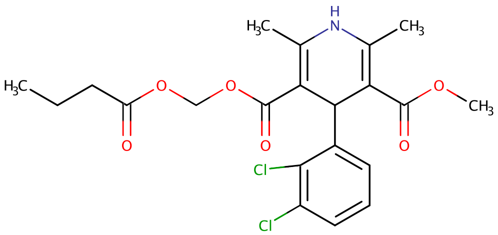

(a) Clevidipine, -9.52 kcal/mol

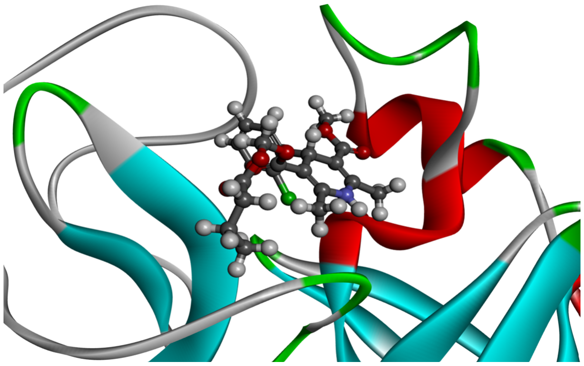

(b) 2019-nCoV protease and Clevidipine complex

Figure 7: Clevidipine and its complex with 2019-nCoV protease.

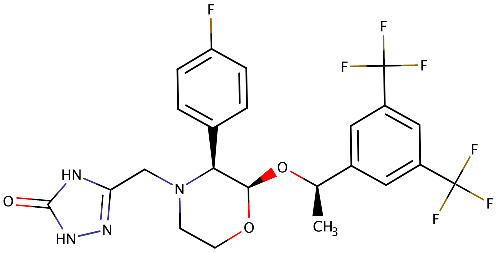

(a) Aprepitant, -9.49 kcal/mol

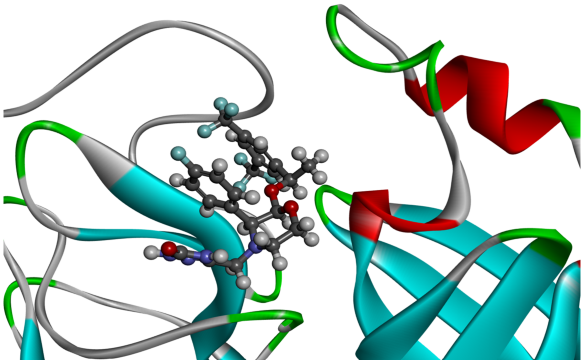

(b) 2019-nCoV protease and Aprepitant complex

Figure 8: Aprepitant and its complex with 2019-nCoV protease.

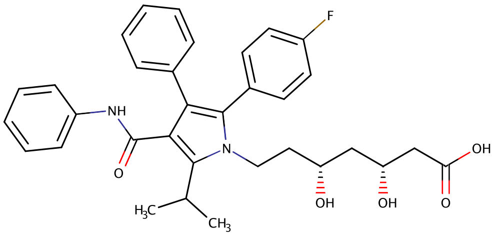

(a) Atorvastatin, -9.49 kcal/mol

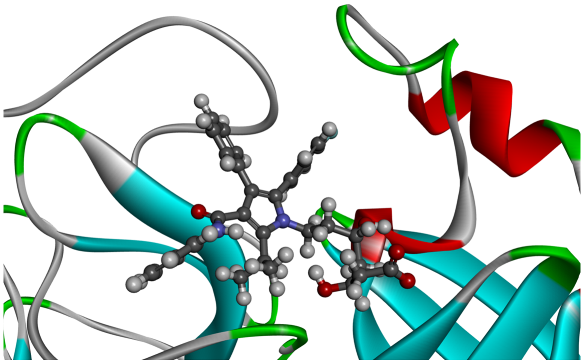

(b) 2019-nCoV protease and Atorvastatin complex

Figure 9: Atorvastatin and its complex with 2019-nCoV protease.

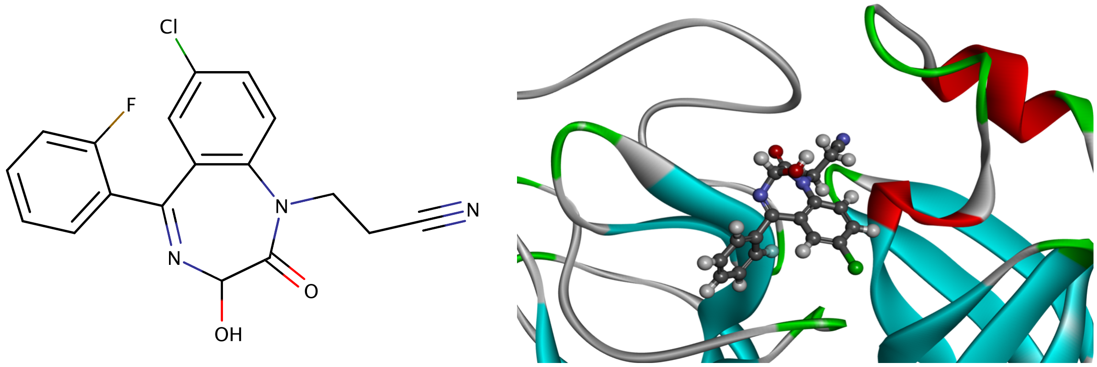

(a) Cinolazepam, -9.47 kcal/mol (b) 2019-nCoV protease and Cinolazepam complex

Figure 10: Cinolazepam and its complex with 2019-nCoV protease.

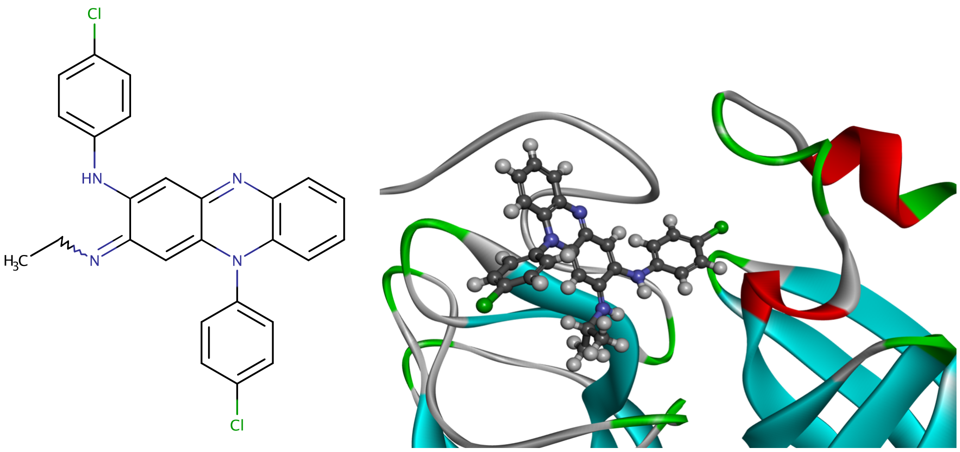

(a) Clofazimine, -9.43 kcal/mol (b) 2019-nCoV protease and Clofazimine complex

Figure 11: Clofazimine and its complex with 2019-nCoV protease.

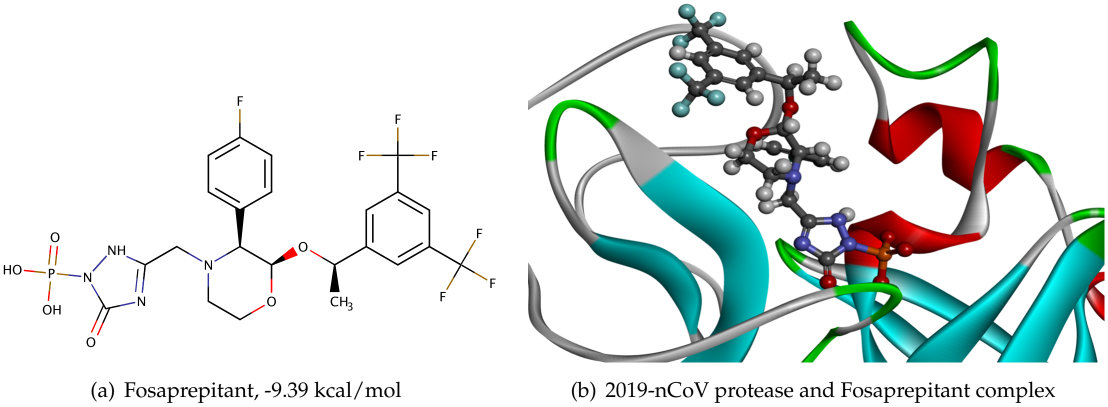

(a) Fosaprepitant, -9.39 kcal/mol (b) 2019-nCoV protease and Fosaprepitant complex

Figure 12: Fosaprepitant and its complex with 2019-nCoV protease.

Figures 2, 3, 4, 5, 6, 7, 8, 9, 10, 11, and 12 display top 4 to 15 molecules predicted by our 3D models. Their predicted binding affinities are given, together with their complexes with 2019-nCoV protease. These compounds are ranked according to their binding affinity values predicted by the consensus of 3DALL and 3DMT.

#### S2 Supplementary Data Guide

Supplementary data are given in TableS1.csv, TableS2.csv, TableS3.csv, FileS2.zip, and FileS1.zip.

**S2.0.0.1 TableS1.csv** Table of the experimental IC50 of 84 SARS-CoV inhibitors.

**S2.0.0.2 TableS2.csv** Table of predicted binding affinities of 1445 FDA-approved drugs and 2019-nCoV protease.

**S2.0.0.3 TableS3.csv** Table of PDBID and experimental affinities of 15,843 complexes in PDBbind v2018 general set.

**S2.0.0.4 FileS1.zip** 3D structures of 84 complexes of SARS-CoV protease inhibitors and 2019-nCoV protease.

**S2.0.0.5 FileS2.zip** 3D structures of 1445 complexes of FDA-approved drugs and 2019-nCoV protease.

**S2.0.0.6 Software** Codes for our deep learning models will be made available to ensure the full reproducibility of the present results.
